# Noradrenergic plasticity of olfactory sensory neuron inputs to the main olfactory bulb

**DOI:** 10.1101/002550

**Authors:** D Eckmeier, SD Shea

## Abstract

Sensory responses are modulated throughout the nervous system by internal factors including attention, experience, and brain state. This is partly due to fluctuations in neuromodulatory input from regions such as the noradrenergic locus coeruleus (LC) in the brainstem. LC activity changes with arousal and modulates sensory processing, cognition and memory. The main olfactory bulb (MOB) is richly targeted by LC fibers and noradrenaline profoundly influences MOB circuitry and odor-guided behavior. Noradrenaline-dependent plasticity affects the output of the MOB. However, it is unclear whether noradrenergic plasticity includes modulation in the glomerular layer, the site of input to the MOB. Noradrenergic terminals are found in the glomerular layer, but noradrenaline receptor activation does not seem to acutely modulate olfactory sensory neuron terminals *in vitro*. We investigated whether noradrenaline induces plasticity at the glomerulus. We used wide-field optical imaging to measure changes in odor responses following electrical stimulation of locus coeruleus in anesthetized mice. Surprisingly, the odor-evoked intrinsic optical signals at the glomerulus were persistently weakened after LC activation. Calcium imaging selectively from olfactory sensory neurons confirmed that this effect was due to a uniform gain suppression of presynaptic input, and did not require exposure to a stimulus during the LC activation. Finally, noradrenaline antagonists prevented glomerular suppression. We conclude that noradrenaline release from LC has persistent effects on odor processing already at the first synapse of the main olfactory system. This mechanism could contribute to arousal-dependent memories.

## Introduction

Neuronal firing patterns are continuously influenced by neuromodulators such as noradrenaline. Noradrenaline is released throughout the forebrain by the brainstem nucleus locus coeruleus (LC) and modulates sensory responses, cognition and behavior with arousal (reviewed in: Usher et al., 1999; Aston-Jones and Cohen, 2005; Bouret and Sara, 2005; Valentino and Van Bockstaele, 2008; Berridge et al., 2012; Devore and Linster, 2012). In the main olfactory bulb (MOB), noradrenaline is essential for certain forms of odor learning and discrimination, including learning of social odors (Pissonnier et al., 1985; Sullivan et al., 1989, 1992, 2000; Kendrick et al., 1991; Rangel and Leon, 1995; Brennan et al., 1998; Guérin et al., 2008; Mandairon et al., 2008). Nonetheless, the specific synaptic targets of persistent modulation by noradrenaline in the MOB remain unclear.

Olfactory sensory neurons (OSNs) make synapses onto mitral/tufted cells which project to deeper brain areas. OSN terminals and the apical dendrites of mitral/tufted cells form spherical glomeruli at the surface of the MOB (Chen and Shepherd, 2005). The glomeruli therefore constitute the first synapse in the main olfactory system. Because the glomeruli gather inputs from OSNs expressing the same receptor protein, they form functional units of spatial representations for odors (Mombaerts, 2006) that may be readily imaged (Rubin and Katz, 1999; Uchida et al., 2000; Meister and Bonhoeffer, 2001; Wachowiak and Cohen, 2001; Bozza et al., 2004; Lin et al., 2006; Fletcher et al., 2009; Ma et al., 2012). Deep to the glomeruli, the mitral/tufted cells receive inhibitory feedback from granule cells via dendrodendritic synapses onto their lateral dendrites (Jahr and Nicoll, 1980; Isaacson and Strowbridge, 1998).

Noradrenaline has both acute and persistent effects on MOB activity. Granule cells are inhibited by α_1_ adrenergic receptor agonists at some concentrations (Nai et al., 2009, 2010; Linster et al., 2011). Mitral/tufted cells are also directly excited by noradrenaline (Hayar et al., 2001; Pandipati et al., 2010). These excitatory and disinhibitory effects may synergize to increase sensitivity to input (Jiang et al., 1996; Ciombor et al., 1999). It is suggested that transient disinhibition of mitral/tufted cells leads to longer-term physiological changes in the MOB network (Pandipati et al., 2010). For instance, the mitral/tufted cells incrementally habituate to an odor presented during noradrenaline release and the response remains suppressed afterwards (Wilson et al., 1987; Sullivan et al., 1989; Shea et al., 2008). This is correlated with a broad increase in levels of GABA relative to glutamate (Kendrick et al., 1992; Brennan et al., 1998) suggesting habituation involves active inhibition.

The circuit nodes affected by this enhanced inhibition remain uncertain. Noradrenaline modulates granule cell activity (Nai et al., 2009, 2010; Linster et al., 2011), and noradrenergic terminals from LC heavily target the granule cells (McLean et al., 1989; Winzer-Serhan et al., 1996, 1997; Day et al., 1997). However, noradrenergic terminals are found throughout the MOB and cells in the glomerular layer express adrenergic receptors (Winzer-Serhan et al., 1996, 1997; Day et al., 1997). These diverse interneurons (many of which are GABAergic) are positioned to regulate synaptic input from OSNs. Activation of noradrenaline receptors failed to acutely modulate signaling from OSNs, but persistent effects were not assessed (Hayar et al., 2001).

We investigated whether persistent modulation of the OSN input to the MOB is induced by electrical LC stimulation during odor presentation in anesthetized mice. We measured the activation of glomeruli with wide-field imaging of intrinsic optical and fluorescent calcium signals at the glomerular layer in the MOB. Surprisingly, after LC stimulation we observed a persistent reduction in signals from the OSNs. This effect was prevented by adrenergic receptor antagonists, but did not require odor-driven activity during LC stimulation. Thus, arousal-dependent olfactory memories may alter odor-evoked synaptic activity as early as the first synapse in the main olfactory system.

## Methods

### Animals

Mice were 6-16 weeks old, of both sexes, and housed in the institution’s animal facilities. Intrinsic optical signals were measured in C57/BL6 mice (Jackson Labs) and fluorescent calcium signals were measured in a transgenic mouse line (tetO-GCaMP2/OMP-IRES-tTA; a gift from CR Yu, Stowers Institute for Medical Research, Kansas City, MO, USA). To generate these mice, a tetO-GCaMP2 mouse line was crossed with another line carrying an OMP-IRES-tTA allele (He et al., 2008; Ma et al., 2012). In this mouse, expression of *OMP* leads to the transcription of a bicistronic RNA for both OMP and tTA (Yu et al., 2004) and in turn to expression of the fluorescent Ca2+ sensor GCaMP2 in olfactory sensory neurons.

### Genotyping (tetO-GCaMP2/OMP-IRES-tTA)

After weaning (d21) tail samples were collected form isofluorane-anaesthetized mice. The samples were solved in lysis buffer (10 mM NaOH and 0.1 mM EDTA) and additional proteinase K at 37°C. The proteinase K was deactivated in a 95°C water bath. The PCR solution included 1 µl of the DNA solution (diluted 20X), 10 µl PCR master mix (Promega GoTaq^®^ Green Master Mix M7123), 7 µl nuclease free water and 1 µl of each primer (10µM; IDENTIFY 5’ and 3’ primers ATCGATTCTAGAATTCGCTGTCTG; CTTATCGTCATCGTCGTACAGAT). <u>PCR cycle</u>: 2 min at 95°C, 35 cycles (30s at 95°C, 1 min 53°C to 59°C, 1 min at 72°C), 5 min at 72°C, stored at 4°C.

### Anesthesia

During all surgeries the mice were initially anaesthetized with ketamine and xylazine (100/5 mg/kg). The anesthesia was maintained with a syringe pump (Harvard Apparatus, Pump 11) loaded with a ketamine/saline solution (i.p. infusion; 90 mg/kg/h).

### Implantation of the Stimulation Electrode

Tungsten electrodes (MicroProbes™; 1MΩ) were shortened to 8 mm from the tip and soldered to new contacts before implantation. Impedence measurement ensured that the electrodes were not damaged. During the procedure the location of the electrode tip was verified through electrophysiology. Locus coeruleus neurons exhibit a characteristic shape and rate of action potentials, and respond to tail pinches (Shea et al., 2008). The electrode was secured with dental cement and the scalp was sutured. The cut was further sealed with tissue adhesive (Vetbond™; n-butyl cyanoacrylate). An anti-inflammatory drug (Meloxicam or Loxicom) was injected to facilitate recovery (1-10 days, 1 mg/kg).

### Acute Cranial Window

A cranial window for imaging was acutely implanted over the MOB immediately before the experiment. A plate with a round window was glued to the exposed skull over the main olfactory bulbs using Super Glue^®^ (cyanoacrylate). After the glue had hardened, the mouse was placed in a warming tube and the head was fixed in a holder via the plate. A craniotomy was opened to reveal the dorsal surfaces of both main olfactory bulbs (compare figures 1 A and 3 A) using a dental drill and a breakable blade holder. Care was taken to remove the bone without injuring the dura mater. The craniotomy was filled with 1.5% agarose (Sigma Aldrich, Agarose Type VII-A, low gelling temperature) and covered with a round (3 mm diameter) cover glass that fits inside the window in the plate. For antagonist experiments, phentolamine (10 µM) and propranolol (10 µM) were dissolved in the agarose used to fill the imaging window. We tested the efficacy of this application method with gabazine and CNQX (see results).

### Imaging Apparatus

The mouse was positioned under a CCD video camera (Vosskühler CCD-1300QF) attached to a macro-lens assembly (Nikon normal AF Nikkor 50 mm auto lens and Nikon telephoto AF DC Nikkor 105 mm lens) (Petzold et al., 2008). The camera was focused on the glomerular layer of the main olfactory bulbs. The aperture was set to 8 for intrinsic optical signals and 2 for fluorescent signals, the focal length was set to infinite. Light emitting diodes (LEDs) of different wavelengths were used for illumination. Intrinsic signals were visible in far/infrared light (∼780 nm), while fluorescence was excited by blue light (470 nm wavelength). A band-pass emission filter was placed in front of the blue LEDs and a 510 nm long-pass optical filter (Chroma Filters HQ510lp) was placed in front of the CCD camera to filter out the blue light from the LED.

### Odor Presentation

The odors were presented using a custom-built olfactometer containing an 8-way solenoid that controls oxygen flow through 8 vials, 7 of which contained odorants dissolved in mineral oil (0.5% or 1%), the remaining vial was empty (blank). Odorized oxygen was diluted 5:1 into a continuous carrier stream for a total flow of 2.5 l/min. To prevent odor accumulation, air was collected behind the animal with a vacuum pump. Odor presentation was 20 s when measuring intrinsic signals and 3 s when measuring calcium signals.

### Electrical Stimulation of Locus Coeruleus

The implanted electrode was connected to an isolated pulse stimulator (A-M Systems Model 2100) and a ground electrode was positioned under the skin behind the mouse’s ear. The pulse generator was triggered 1 s prior to odor presentation onset. The stimulation consisted of 40 μA biphasic pulses of 100 μs duration generated at 5 Hz for 24 s (intrinsic optical imaging) or 5 s (calcium imaging).

### Automated Experimental Control and Data Acquisition

Custom software written in LabView (written by DE) controlled the data acquisition hardware and stimulus delivery according to a scripted protocol. The olfactometer, LEDs, and pulse stimulator were controlled by a digital I/O card (National Instruments, PCI-6503) connected with the devices via a terminal block (National Instruments, CB-50LP). For camera control and image acquisition a separate image acquisition card (National Instruments IMAQ PCI-1422-LVDS).

### Imaging Protocol

During each trial, images were acquired from the camera before, during and after odor presentation and stored as monochromatic image stacks. For intrinsic signals we acquired 250 frames before, 500 frames during and another 250 frames after odor presentation at 25 frames per second (fps). For fluorescent signals we acquired 100 frames before, 75 frames during and 50 frames after odor presentation at 25 frames per second (fps). Differences in the frame counts for the two measurements reflect the different time scales necessary to acquire reliable signals.

### Experimental Sequence

First, 7 different odors and clean air were each presented three times in a ‘pretest’. Based on the responses, two odors were selected for the experiment. The two odors were presented alternately throughout the experiment with an interstimulus interval of 70 s (30 s for the moderated protocol.. Experiments had a pairing phase of 30 repetitions during which both odors were presented and one of which was paired with locus coeruleus stimulation. Imaging phases span 15–20 repetitions and occurred before and after the pairing phase. In the results, an experiment is defined as performance of the above procedure in one mouse and yielded measurement of two odor responses from either one or two hemispheres (ipsilateral and/or contralateral). Each mouse was stimulated only once.

### Verification of the Stimulation Site

After the experiment ended, the position of the stimulation electrode was marked by an electrolytic lesion using a pulse stimulator (three pulses at −10 μA, 10 s). Then the mouse was given a lethal injection of Euthasol and perfused with PBS followed by 4% paraformaldehyde. The skull was stored in paraformaldehyde overnight before the brain was extracted. The brain was then placed in 30% sucrose in PBS overnight, cut into 60μm slices and stained with cresyl violet.

### Data analysis

For quantification, the raw image stacks were analyzed with our custom Matlab (Mathworks, RRID:nlx_153890) software (written by DE). Active areas on the olfactory bulb were made visible by dividing the average frame image acquired during stimulation by the average frame image acquired prior to stimulation (Meister and Bonhoeffer, 2001). Typical results are shown in Figures 1A and 2A. Regions of interest that corresponded to activated glomeruli for each odor were identified and the mean grayscale value was computed frame-by-frame from the raw image data. This produced a signal trace for every repetition and every glomerulus. The single repetition traces were then averaged for repetitions before and after odor pairing, respectively. Slow baseline shifts were corrected by fitting an exponential function to the baseline and subtracting it from the data. This analysis resulted in traces like those shown in Figures 1B and 2B. Only glomeruli that showed a clear response correlated with odor onset in at least one of the experimental phases were included in the data set. From the traces we calculated the signal strength for each experimental phase as (|Rb-Ra|)/Rb*100 with Rb being the mean baseline reflectance/fluorescence and Ra the mean reflectance/fluorescence during odor stimulation (plateau phase).

**Figure 1.**
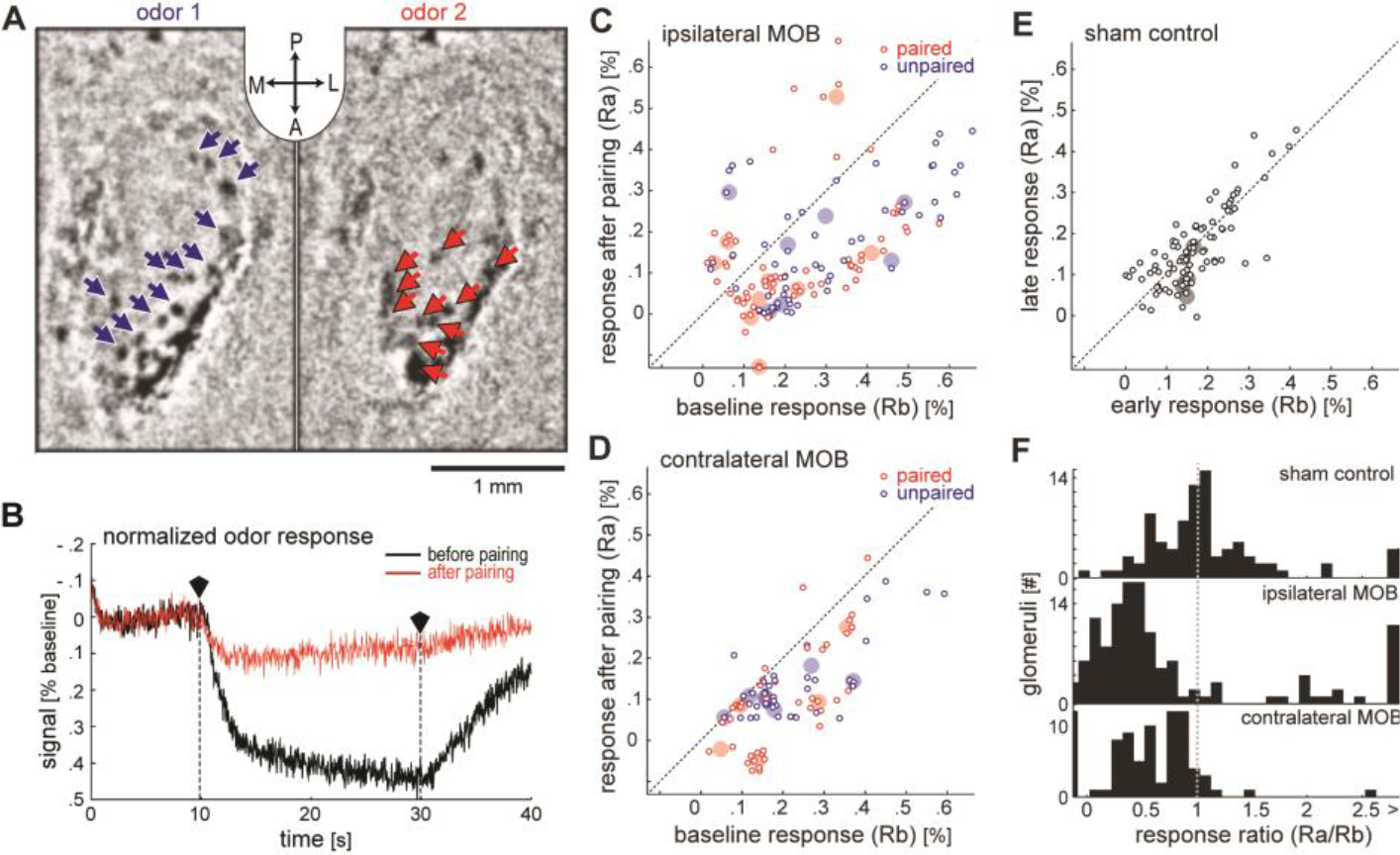
Intrinsic optical signals reveal suppression of glomerular responses after pairing a series of odor presentations with locus coeruleus stimulation. ***A***, The response of glomeruli on the dorsal surface of the left hemisphere of the MOB to two odors as imaged with IOS (inset: A-anterior, P-posterior, L-left, R-right). The blue and red arrows denote each of the responsive glomeruli identified by the analysis. These images were generated by subtracting images acquired during odor presentation from those acquired prior to odor presentation. ***B***, An average response before locus coeruleus stimulation (black trace) is contrasted with the average response trace afterwards (red trace) in the same glomerulus. The two traces indicated illumination over time. The traces were each averaged over 20 rials, corrected for a slow baseline decay (see Methods), and then normalized to their respective average value before odor onset. The Y axis therefore indicates the deviation from baseline activation in percent of the baseline. We inverted the Y axis so that higher values indicate stronger activation. ***C-E***, The Y axis indicates response strength after pairing while the X axis indicates response strength before pairing. Small circles indicate the activity of glomeruli, and larger filled circles indicate the median glomerular response for each animal and odor. C and D show the data for glomeruli responding to the paired odor (red) and unpaired odor (blue) on the hemisphere ipsilateral and contralateral to the stimulated locus coeruleus, respectively (9 animals). Panel E shows the results of the sham control (4 animals). Data from both odors and hemispheres were combined in this case. ***F***, The ratio between the response before and after pairing was calculated for each odor. The histograms display the distribution of glomeruli for the sham control, and for the actual experiment divided by hemisphere.

**Figure 2.**
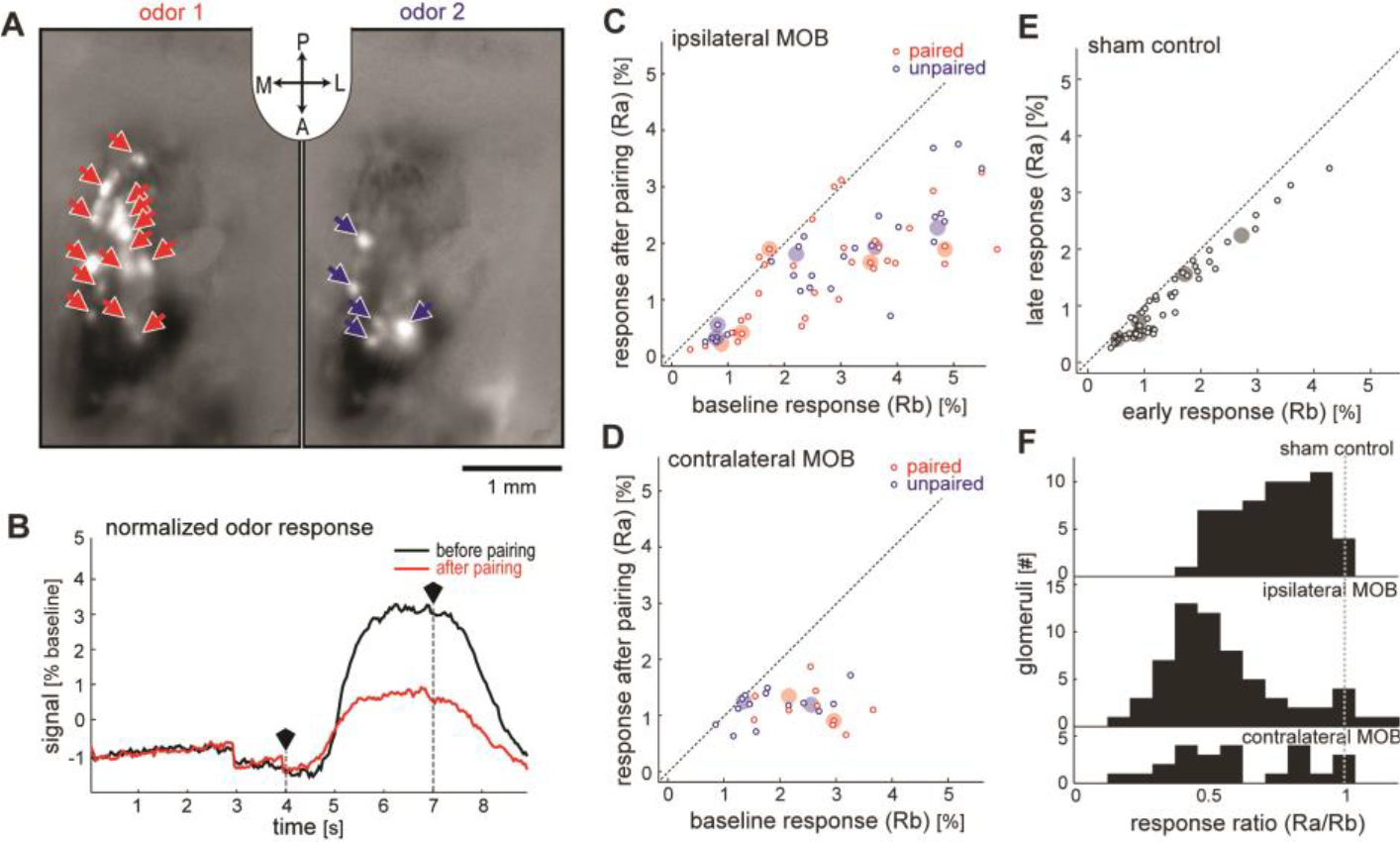
Fluorescent OSN calcium signals reveal suppression of glomerular responses after pairing a series of odor presentations with locus coeruleus stimulation. ***A***, The response of OSNs terminals (glomeruli) on the dorsal surface of the left hemisphere of the MOB to two odors as imaged with GCaMP2 (inset: A-anterior, P-posterior, L-left, R-right). The blue and red arrows denote each of the responsive glomeruli identified by the analysis. These images were generated by subtracting images acquired during odor presentation from those acquired prior to odor presentation. ***B***, An average response before locus coeruleus stimulation (black trace) is contrasted with the average response trace afterwards (red trace) in the same glomerulus. The two traces indicated illumination over time. The traces were each averaged over 20 trials, corrected for a slow baseline decay (see Methods), and then normalized to their respective average value before odor onset. The polygonal black markers and vertical dotted lines mark beginning and end of odor presentation. Due to the normalization, the Y axis indicate the deviation from baseline activation in percent of the baseline. ***C-E***, The Y axis indicates response strength after pairing while the X axis indicates response strength before pairing. Small circles indicate the activity of glomeruli, and larger filled circles indicate the median glomerular response for each animal and odor. ***C and D*** show the data for glomeruli responding to the paired odor (red) and unpaired odor (blue) on the hemisphere ipsilateral and contralateral to the stimulated locus coeruleus, respectively (5 animals). Panel ***E*** shows the results of the sham control (5 animals). Data from both odors and hemispheres were combined in this case. ***F***, The ratio between the response before and after pairing was calculated for each odor. The histogram indicates the distribution of glomeruli for the sham control, and for the actual experiment divided by hemisphere.

### Statistics

The Kruskal-Wallis test (a non-parametric test for differences between independent distributions) followed by Tukey’s honestly significant difference test was used to identify statistically significant differences (Matlab, Mathworks).

## Results

We aimed to assess the persistent effects of noradrenaline from LC on odor responses in the glomeruli. We chose to use wide-field imaging because it allowed us to measure the amplitude of glomerular signals over an extended time period for multiple glomeruli (Rubin and Katz, 1999; Uchida et al., 2000; Meister and Bonhoeffer, 2001; Wachowiak and Cohen, 2001; Bozza et al., 2004; Lin et al., 2006; Fletcher et al., 2009; Ma et al., 2012). As an initial step, we imaged intrinsic optical signals (IOS), which reveal spots of activity that correspond to glomeruli.

### IOS are suppressed after pairing odors with LC stimulation

We measured the effects of LC activation on IOS in the MOB. Figure 1A shows two differential images (mean odor frame minus mean baseline frame) of the left MOB. Images were processed as described in Methods. Dark spots represent regions showing a change in reflectance of infrared light (indicative of neural activity) in response to two odors. Arrows indicate the locations of all activated glomeruli identified by our analysis in this example. Figure 1B shows an example of the temporal profile of the signal observed in one of these glomeruli. There is a clear deflection of the signal just after odor onset and a return to baseline after odor offset. As in this example, substantial attenuation of the signal was typically observed following LC stimulation paired to the odor. To quantify this result, we paired the presentation of one of two odors with electrical stimulation of LC via an implanted electrode thirty times. Before and after stimulation we imaged the response of the glomeruli activated by both odors for 20 trials each. Sham control mice were treated identically, but we did not apply stimulation current to LC.

In mice that received no stimulation current in LC (‘sham’ condition), the glomerular IOS measured before and after LC stimulation were indistinguishable. In Figure 1E, the mean response strength for the last 20 trials is plotted over the mean response strength for the first 20 trials, for each glomerulus (n = 88, from both hemispheres; n = 4 mice). The large transparent dots indicate median values for each animal. Clustering of the data along the unity line demonstrates that response strengths were stable and unaffected by sham LC stimulation.

In contrast, in mice that did receive LC stimulation in conjunction with odors (‘paired’ condition), IOS signals in response to both odors were largely suppressed from baseline levels during the post-stimulation phase (Figure 1C, D). Most glomeruli on both hemispheres exhibited reduced responses after the pairing (n = 127 glomeruli, ipsilateral MOB; n = 82 glomeruli, contralateral MOB; n = 9 mice), as the data points are mostly below the unity line. In some cases, the suppression was so complete that imperfect baseline correction of unresponsive glomeruli left a small negative signal. Note also that some glomeruli increased responses after the pairing, but almost all of them derived from two mice tested on separate occasions. One of these mice yielded responsive glomeruli on the contralateral hemisphere that were all suppressed. In light of these observations, we are hesitant to conclude that enhancement is a widespread occurrence.

Figure 2F shows histograms of the ratio between the post-stimulation and baseline response strengths (suppression ratio, see Materials and Methods) for all glomeruli in control mice, ipsilateral glomeruli in paired mice, and contralateral glomeruli in paired mice. For both populations of glomeruli (ipsilateral and contralateral) in paired mice, this ratio was significantly lower than 1 (for the non-normally distributed data we report the median and the upper bounds of the 1^st^ and 3^rd^ quartiles [Q_1_ and Q_3_ respectively]. ipsilateral: median: 0.44, Q_1_: 0.25, Q_3_: 0.76; contralateral: median: 0.58, Q_1_: 0.30, Q_3_: 0.84; Wilcoxon signed rank test, contralateral: p = 0.000, ipsilateral: p = 0.000), However this was not the case for glomeruli in control mice (median: 0.9975, Q_1_: 0.728, Q_3_: 1.27; Wilcoxon signed rank test, p = 0.89). Comparison of the three distributions showed that both paired distributions significantly differed from the control distribution (Kruskal-Wallis, χ ^2^ = 53.75, p = 0.000, posthoc Tukey’s HSD) but not from each other (posthoc Tukey’s HSD).

Although only one of the odors was paired with LC stimulation, the suppression of glomerular responses was observed for both odors. Comparison between the distributions of suppression ratio for the paired odor and the unpaired odor yielded no significant difference (Kruskal-Wallis, χ ^2^ = 0.86, p = 0.35). Thus, LC-mediated plasticity of glomerular responses is not odor specific for this simulation protocol.

### Fluorescent calcium signals in OSNs are suppressed after pairing odors with LC stimulation

While intrinsic optical signals are closely correlated with the activity of OSNs, the extent of contributions from other glomerular elements to these signals remains unclear. We hypothesized that the changes we observed in glomerular activity after LC stimulation were due to reduced synaptic input from OSNs. Therefore, we moved on to more directly measure OSN synaptic input using wide-field fluorescence imaging in a mouse line that expresses the genetically encoded calcium sensor GCaMP2 exclusively in OSNs (OMP-GCaMP2) (Yu et al., 2004; He et al., 2008; Ma et al., 2012).

We treated raw data for GCaMP2 fluorescence signals from OSNs in the same manner as data from IOS experiments (Figure 2A, B). Figure 2A shows two differential images of the left MOB that reveal changes in GCaMP2 fluorescence evoked by two different odors. The round bright dots are glomeruli that were activated in response to the stimulus. Arrows indicate responsive glomeruli that were identified and selected for further analysis. As with the IOS data, we corrected for slow baseline decay in brightness. Comparison of the mean responses taken from a representative glomerulus before and after the stimulation phase reveals a clear response suppression after LC stimulation (Figure 2B).

In these experiments, we measured the effects of LC stimulation specifically on the presynaptic input from OSNs in OMP-GCaMP2 mice. As before, we repeatedly paired the stimulation of LC with the presentation of one of two odors and imaged the response to both odors before and after this stimulation phase. Figure 2C-F depicts the results of these experiments in the identical format as in Figure 1C-F. In the absence of LC stimulation (‘sham’ condition), the calcium signals from OSNs at the glomeruli did not change (n = 58, from both hemispheres; n = 5 mice). This is illustrated by the scatter plot in Figure 2E, in which the data for most glomeruli fall close to the unity line.

However as seen with IOS, the fluorescent calcium response to odors was suppressed after LC stimulation (Figure 2C,D). The measurements for nearly all glomeruli on both hemispheres (n = 62 glomeruli, ipsilateral MOB; n = 24 glomeruli, contralateral MOB; n = 5 mice) were at or below the unity line. Plotted as histograms of suppression ratio, the distributions for ipsilateral and contralateral glomeruli in paired mice were significantly lower than 1 (ipsilateral: median: 0.52, Q_1_: 0.41, Q_3_: 0.66; contralateral: median: 0.54, Q_1_: 0.42, Q_3_: 0.83; Wilcoxon signed rank test, contralateral: p = 0.000, ipsilateral: p = 0.000). In contrast to the findings with IOS, the sham control mice also show a deviation from 1 which we attribute to bleaching (median: 0.74, Q_1_: 0.62, Q_3_: 0.88; Wilcoxon signed rank test, p = 0.000). Comparison of the populations shows that the distributions of ipsilateral and contralateral glomeruli differ significantly from the distribution of control glomeruli (Kruskal-Wallis; χ^2^ = 25.3; p = 0.000; posthoc Tukey’s HSD), but not from each other.

As was the case with the IOS data, the suppression of glomerular responses was evident for both the paired and unpaired odors. Comparing the suppression ratios for glomeruli responsive to the paired and the unpaired odor, we found there was not a significant difference (Kruskal-Wallis; χ^2^ = 3.02; p = 0.083). These data confirm that LC-mediated plasticity of OSN input to the glomeruli is not odor specific for this pairing regime.

Further analysis of the changes across all glomeruli are consistent with LC stimulation causing a uniform change in the gain of presynaptic input to the MOB. In other words, the degree of suppression was the same regardless of the initial strength of the signal in a given glomerulus. Figure 3A shows that the data from Figure 2C are well fitted to a linear function that passes through the origin. For the ipsilateral paired glomeruli, responses seen after pairing were significantly linearly correlated to those seen at baseline (Pearson correlation coefficient *r* = 0.79, *p* = 0.000). Figure 3B further shows that the magnitude of suppression for glomeruli was unrelated to their baseline response strength (Pearson correlation coefficient *r* = −0.15, *p* = 0.16). These observations are consistent with LC stimulation pairing uniformly scaling input to all active glomeruli. As shown in Figure 3A (inset), this modulation was not evident in unresponsive glomeruli. Our data further reveal that this effect is not simply due to changes in resting fluorescence. No change in the absolute brightness of responsive glomeruli during baseline or odor periods was observed in sham controls (Figure 3C). While there was evidence of absolute brightness changes after LC stimulation, those changes were comparable during baseline and odor periods (Figure 3D).

**Figure 3.**
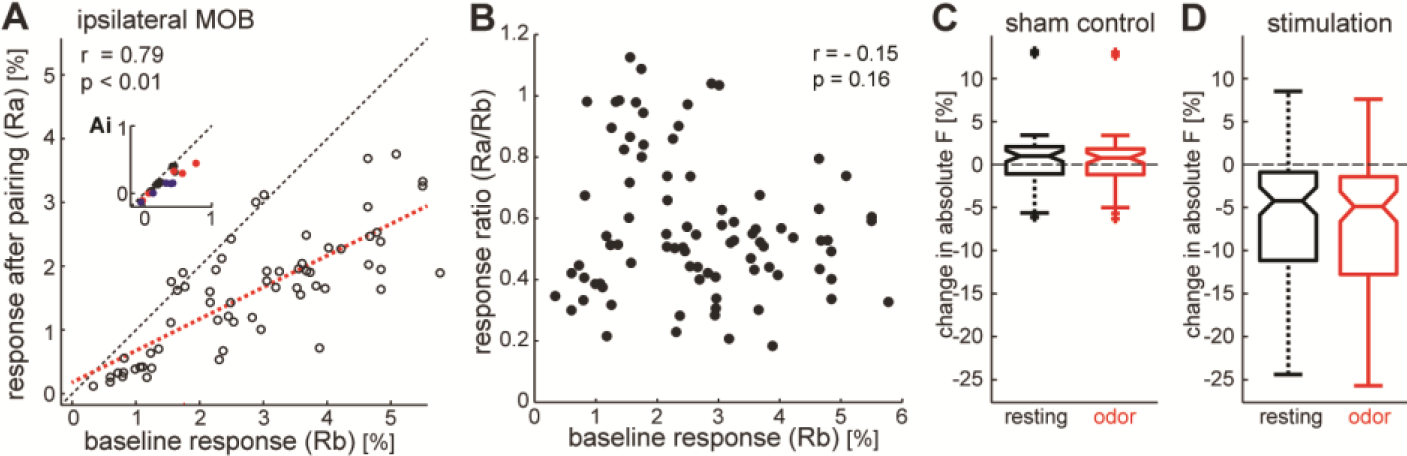
The pattern of suppression across glomeruli is consistent with a divisive gain mechanism. ***A***, A plot of all ipsilateral glomerular responses measured with GCaMP2 imaging from Figure 2C, plotted with the same conventions. Here, glomeruli responding to the paired and unpaired odors are not distinguished. Note that baseline and post-pairing responses are linearly correlated and that the scatterplot is fit with a line that passes through the origin. The inset shows a comparable plot (at the same scale) of mean activity during the odor among non-responsive regions of the MOB. ***B***, A scatterplot of suppression ratio as a function of the amplitude of baseline responses for each glomerulus. The two variables are not significantly correlated. ***C and D***, Box plots of the distributions of changes in absolute fluorescence seen over the glomeruli plotted in A and B following sham stimulation (C) and LC stimulation (D). In each plot, changes in absolute fluorescence are indicated for the pre-odor (‘resting’) period (black) and the odor period (red).

Odor responses imaged in OMP-GCaMP2 mice were sufficiently strong and reliable to permit examination of the fluctuations of responses from trial to trial. In figure 4A-D we show the trial by trial data for these experiments. The scatter plots indicate the response strength of each glomerulus normalized to the baseline activity during each odor presentation. We did not image during the LC stimulation period (repetitions 21-50) to limit bleaching effects. A general downward shift is apparent, however glomeruli which were activated during LC stimulation were further suppressed.

**Figure 4.**
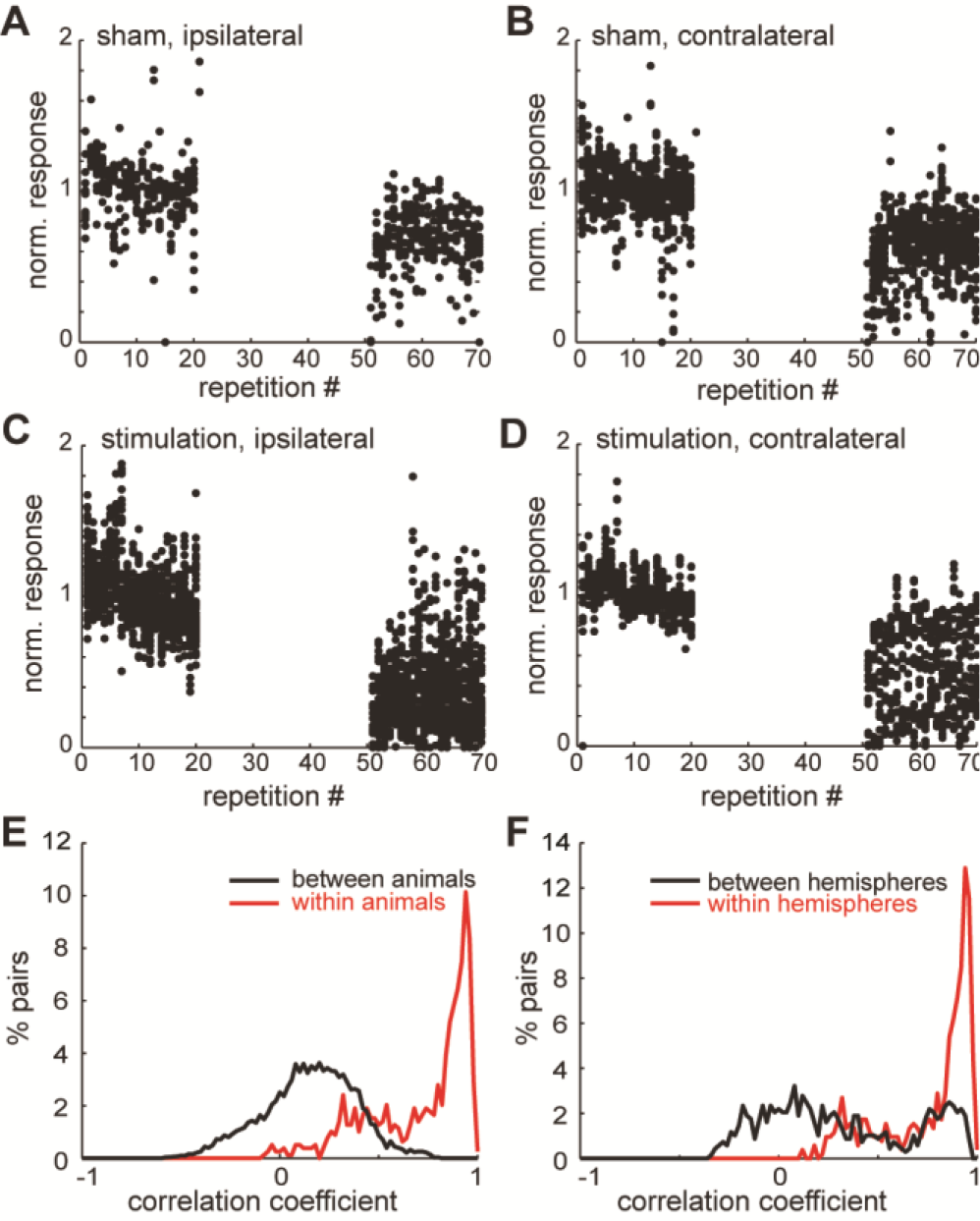
Trial by trial calcium signals and correlation between glomeruli. We show the calcium response of all glomeruli to the two odors on each hemisphere from all animals. All plots indicate the response strength during the experiment, normalized to the mean activation during the first 20 odor presentations, as the baseline varied for each glomerulus. No imaging was performed during the locus coeruleus activation. ***A and B***, For sham control experiments, the responses of glomeruli on the hemispheres ipsi- and contralateral to the stimulation electrode, respectively. ***C and D***, Corresponding data for animals undergoing LC stimulation. ***E and F***, Correlation histograms of percent pairs over correlation coefficient (bin size 0.04). Autocorrelations were omitted. In E the red line corresponds to pairs of glomeruli responding to the same odor in the same mouse. The black line shows correlation of glomeruli from different animals. In F the red line corresponds to pairs of glomeruli which respond to the same odor and are located in the same hemisphere. The black line corresponds to pairs of glomeruli which respond to the same odor but were found on different hemispheres of the MOB.

The change in signal strength over repetitions in glomeruli which respond to the same odor and are located on the same hemisphere were highly correlated. To show correlations between groups of glomeruli we present correlation histograms as percent of pairs over the correlation coefficient (bin size = 0.04 bins) in Figure 4E, F. In Figure 4E we show that glomeruli which respond to the same odor within a mouse correlate well (red line) while glomeruli do not correlate across animals (black line). In Figure 4 F we show that glomeruli which respond to the same odor but are localized on different hemispheres correlate only weakly (black line) while those which are located on the same hemisphere correlate well.

### Effects of LC stimulation on OSNs depend on noradrenaline receptors

Using two independent imaging methods we showed that LC stimulation triggers persistent changes in glomerular responses to odors, and we attribute these changes to reduced synaptic activity at OSN terminals. The question remains whether these effects are caused by noradrenaline release in the glomerular layer, or they involve an indirect mechanism. To answer this question, we topically applied the α- and β-noradrenergic receptor antagonists phentolamine and propranolol to the MOB surface during imaging experiments.

Drugs were dissolved in the agarose that was placed under the cranial window (see Materials and Methods). We performed electrophysiology control experiments with gabazine (10 µM; n = 2) and NBQX (10 µM; n = 2) to validate this delivery method. Gabazine and NBQX reliably and reversibly excited and suppressed activity respectively in the mitral cell layer (Figure 5) when dissolved in agarose and applied to the dorsal surface of the MOB. In pilot experiments, we observed that medium - high doses (> 60µM) of phentolamine and propranolol alone or in combination abolished odor responses (data not shown). When we lowered the concentration to 10 µM, the odor responses remained stable throughout a timeframe suitable for our experiments. Given the low dose of antagonists, and in order to operate within the potentially limited window of drug effects, we chose to apply a moderated LC stimulation protocol as described by (Shea et al., 2008).

**Figure 5.**
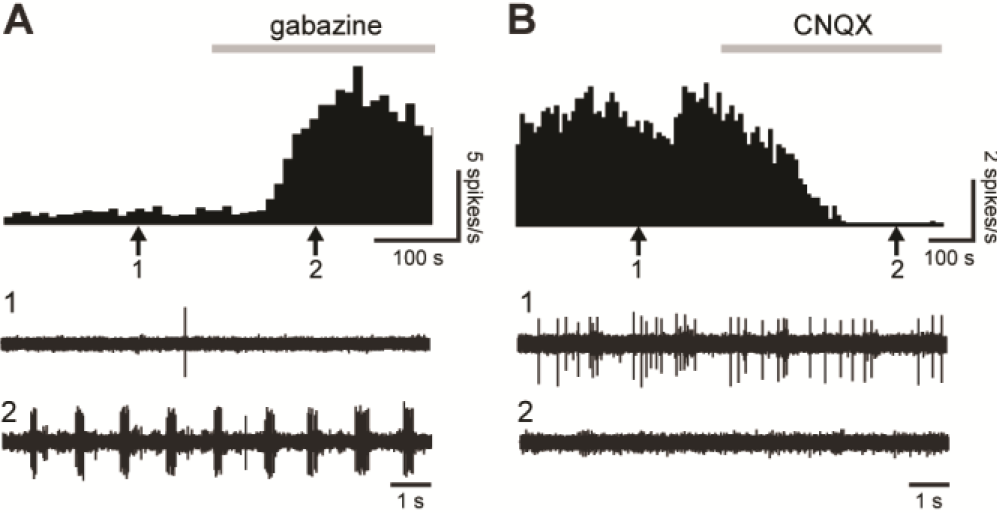
Application of drugs in agarose to the MOB surface effectively manipulates glomerular transmission. We dissolved gabazine (A) or CNQX (B) in agarose (10 µM) and applied the agarose to the surface of the MOB. Mitral cell firing was measured with extracellular electrophysiology. ***A and B***, The upper panel shows a histogram (bin size: 10s) of spike rates during application of gabazine (A) or CNQX (B). The gray bar indicates the application of drug. The middle (1) and bottom (2) panels show raw recording traces taken at time points corresponding with the arrows 1 and 2 respectively. Gabazine successfully disinhibited mitral cells and CNQX successfully inhibited mitral cell firing.

In the absence of antagonists, this LC stimulation regime reliably resulted in a suppression of fluorescent calcium signals after the stimulation phase (Figure 6A; n = 48 glomeruli; n = 3 mice). The magnitude of this suppression was significantly attenuated in the presence of phentolamine and propranolol (Figure 6B; n = 48 glomeruli; n = 3 mice). Post-pairing responses were significantly linearly related to the strength of baseline responses in control (Pearson correlation coefficient *r* = 0.93, *p* = 0.000) and NA antagonist (Pearson correlation coefficient *r* = 0.87, *p* = 0.000) experiments (Figure 6A,B). The histogram of suppression ratio in Figure 6C reveals a rightward shift in the distribution with antagonists (median: 0.81, Q_1_: 0.71, Q_3_: 0.88; Wilcoxon signed rank test, p = 0.000) as compared to the distribution of glomeruli without antagonists (median: 0.60, Q_1_: 0.53, Q_3_: 0.68; Wilcoxon signed rank test, p = 0.000). These two distributions were significantly different (Kruskal-Wallis, χ ^2^ = 37.48, p = 0.000). Thus, the observed suppression of OSN input to the MOB after LC stimulation at least partly depends on noradrenaline receptor activation in the superficial MOB.

**Figure 6.**
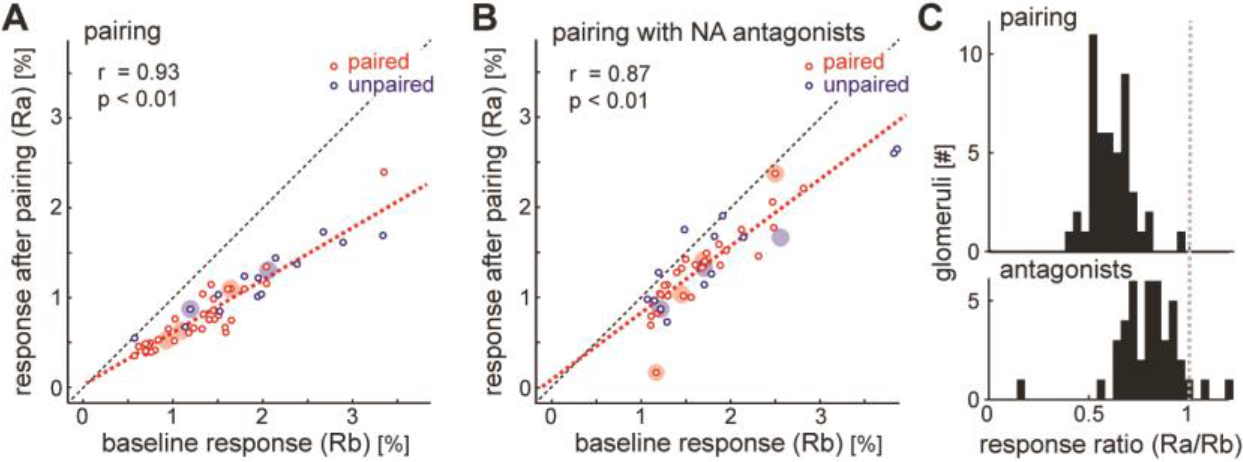
Blocking noradrenalin receptors abolished the suppression of glomerular responses observed after LC stimulatiom. ***A and B***, The Y axis indicates response strength after pairing while the X axis indicates response strength before pairing. Small circles indicate the activation of glomeruli, larger filled circles indicate the median for each animal and odor. A and B show glomeruli responding to the paired odor (red) and unpaired odor (blue) on both hemispheres, for the animals treated with noradrenalin antagonists (B, 4 animals) and saline control animals (A, 3 animals). ***C***, The ratio between the response before and after pairing was calculated for the glomeruli in each group and the distribution is illustrated in the histogram.

### Effects of LC stimulation on OSNs do not depend on odor activity

Since odor responses were suppressed for both paired and unpaired odors and suppression was therefore non-specific, we directly tested whether LC stimulation could suppress glomerular responses in the absence of a paired odor (‘unpaired LC stimulation’). These experiments were performed in the same manner as the moderated pairing protocol above with the exception that no odor was presented during the stimulation phase. Remarkably, in contrast to the results with this protocol described above, unpaired LC stimulation dramatically and invariably suppressed all responsive glomeruli in all experiments (Figure 7A). The close linear relationship between baseline and post-stimulation responses (Pearson correlation coefficient *r* = 0.93, *p* = 0.000) was again consistent with uniform scaling of responses to odors not presented during the stimulation phase. The histograms of suppression ratio in Figure 7B reveal a leftward shift in the distribution from unpaired LC stimulation (median: 0.44, Q_1_: 0.38, Q_3_: 0.47; Wilcoxon signed rank test, p = 0.000) as compared to that of not only sham stimulation, but also paired LC stimulation. All three distributions were significantly different (Kruskal-Wallis, χ ^2^ = 104.3, p = 0.000). Therefore, LC suppression was actually stronger when stimulation was performed in the absence of an odor.

**Figure 7.**
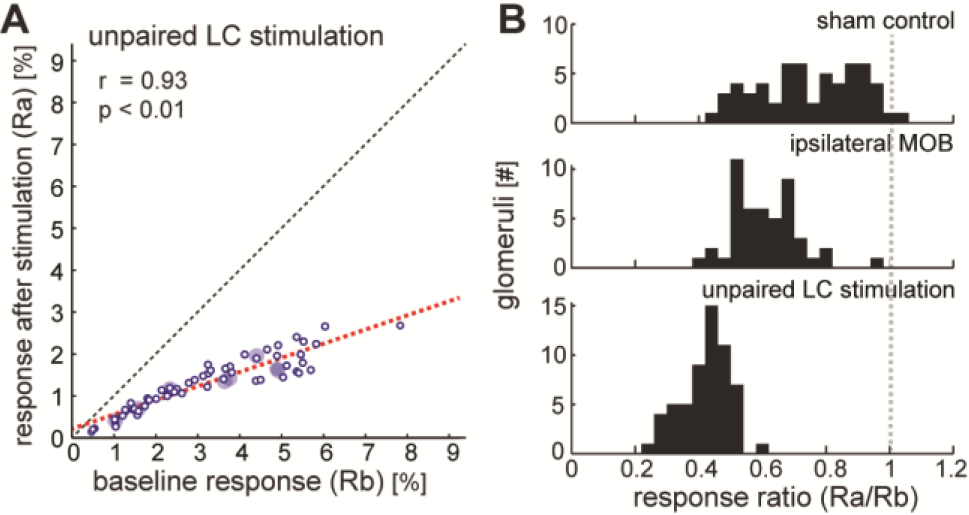
The effects of LC stimulation on glomerular responses did not require odor activity during stimulation. ***A***, The Y axis indicates response strength after unpaired LC stimulation (stimulation in the absence of an odor) while the X axis indicates response strength before stimulation. Small circles indicate the activation of glomeruli, large filled circles indicate the median for each animal odors (4 animals). The plot shows glomeruli on both hemispheres responding to either of two odors. ***B***, The ratio between the response before and after unpaired stimulation was calculated for all glomeruli and the distribution is illustrated in the histogram. For comparison, the identical distributions are shown for sham control experiments (top) and LC stimulation using the same protocol that was paired to an odor (middle).

## Discussion

Here we investigated noradrenergic modulation of synaptic input from olfactory sensory neurons to the glomeruli of the main olfactory bulb in anesthetized mice. By comparing the magnitude of IOS odor responses in the glomeruli before and after LC stimulation during odor exposure, we found that glomerular activity was suppressed long-term by noradrenaline. Subsequent imaging experiments using mice expressing the genetically-encoded calcium sensor GCaMP2 selectively in OSNs allowed us to confirm that this suppression is implemented as a reduction in the gain of presynaptic input to the glomeruli. Pharmacological manipulations revealed that LC-mediated suppression required noradrenergic receptor activation in the glomerular layer. Exposure to an odor during stimulation is however, not required. Instead, locus coeruleus stimulation in the absence of odors resulted in an even stronger reduction of odor response.

We were surprised to find that OSN terminals undergo persistent modulation by noradrenaline. Although noradrenergic fibers from LC course through all layers of the MOB (McLean et al., 1989), and noradrenergic receptors are present in the glomerular layer (Winzer-Serhan et al., 1996, 1997; Day et al., 1997), these inputs are sparser than those to deeper layers. Moreover, an *in vitro* study that examined modulation of signaling in the glomerulus did not detect any acute changes in the presence of noradrenaline (Hayar et al., 2001). Interestingly, odor conditioned fear memories were also recently shown to modify presynaptic glomerular inputs (Kass et al., 2013).

The role of granule cells in the noradrenergic modulation of mitral/tufted cells has been the subject of more intensive study than the glomerular processes. *In vitro* studies of the MOB circuit have established that noradrenaline acutely enhances the sensitivity of mitral/tufted cells to odor input directly and via release of inhibition from granule cells (Hayar et al., 2001; Nai et al., 2010; Pandipati et al., 2010; Linster et al., 2011). These *in vitro* observations are consistent with the acute effects of LC stimulation *in vivo* (Jiang et al., 1996). The synergy of sensitization and disinhibition of the mitral cells has been proposed as a key step in initiating long-term changes to the MOB network, including habituation or suppression of mitral/tufted cells and enhancement of oscillatory rhythms (Brennan and Keverne, 1997; Gire and Schoppa, 2008; Shea et al., 2008; Pandipati et al., 2010).

Shea et al (2008) also stimulated LC *in vivo* during odor presentation and found that odor responses in mitral/tufted cells were suppressed after LC-odor pairing. The effects we see on OSNs appear very similar. It seems likely that the reduced odor response in the mitral/tufted cells is, at least in part, a consequence of the effects we found in the olfactory sensory neurons. We also stimulated LC unilaterally; however by imaging both olfactory bulbs we observed OSN input suppression in both hemispheres of the main olfactory bulb. This was not completely unexpected as it is known that a minority of noradrenergic LC neurons decussate and terminate in the contralateral MOB. Consistent with this asymmetry, the suppression in the contralateral hemisphere was smaller than in the ipsilateral hemisphere.

Our data contrast with the results of the previous study in mitral/tufted cells (Shea et al., 2008) in one important respect. Although we presented two odors during the pairing phase and paired only one of them with LC stimulation, significant suppression of glomerular responses was observed for both odors as compared to sham stimulation experiments. There was a tendency for the suppression of responses to the unpaired odor to be weaker than the suppression of responses to the paired odor in GCaMP2 imaging experiments using 20 s LC stimulation trains, however this difference was not statistically significant (p = 0.083). A weak bias such as this could conceivably be amplified among the mitral/tufted cells through additional modulation by granule cells.

How does plasticity of OSN input to the MOB occur? One possibility is that OSN synaptic terminals undergo enhanced inhibition from periglomerular cells. Periglomerular cells are GABAergic and regulate synaptic input to the glomeruli through presynaptic inhibition of OSN terminals (Murphy et al., 2005). Thus, coordinated modulation of this network could plausibly implement the uniform gain suppression we see here. These cells could also be a source for increased GABA release in response to a memorized odor that is observed *in vivo* after noradrenaline-dependent odor learning (Kendrick et al., 1992; Brennan et al., 1998). However, further detailed study will be required to test this speculation.

At first, it may seem counterintuitive that sensory responses to a learned stimulus would be reduced rather than sensitized. However, habituation - diminishing behavioral responses to a repeated and therefore familiar stimulus - is one of the most fundamental forms of learning. Behavioral habituation is frequently manifested in the brain by neuronal habituation (Horn, 1986). Moreover, sensory plasticity frequently includes dynamic increases in inhibition (reviewed in: Carcea and Froemke, 2013). Broad inhibition is an important component for enhancing the salience or ‘signal-to-noise ratio’ by filtering redundant or overlapping portions of competing representations (Olshausen and Field, 2004; Assisi et al., 2007; Koulakov and Rinberg, 2011; Sachdev et al., 2012; King et al., 2013). We speculate that this may be the case here. Indeed, there is some indication in our data that responses to odors presented during the pairing phase may be partially protected from even stronger suppression. Such a bias could enhance sparseness. Ultimately, ‘sparsening’ of the representation will entail increases in the activity of a small number of neurons in response to the learned stimulus against a background of global response reduction. This may be implemented as a two-stage process consisting of a nonselective subtraction and rectification step followed by multiplicative amplification. We speculate that noradrenergic modulation of the glomeruli may be a mechanism for implementing the first stage of representational sparsening.

Putting our findings into a behavioral context, several forms of noradrenaline-dependent memories are accompanied by physiological changes in the main olfactory bulbs (Wilson et al., 1987; Sullivan et al., 1989; Yuan et al., 2002). Furthermore, selective behavioral and physiological changes in response to an odor can be induced by increasing release of noradrenaline in the presence of that odor (Sullivan et al., 2000; Shea et al., 2008). This simulates what would occur during an episode of arousal. We therefore argue that there is strong evidence that LC-mediated plasticity in the olfactory bulb constitutes an important mechanism for arousal to facilitate odor memory formation. Surprisingly, these memories seem to affect even the initial detection of a stimulus by altering the signal as early as in the receptor neurons.

